# Impact of carbon monoxide on neural activation during a reaction time task

**DOI:** 10.1101/2023.01.17.524443

**Authors:** Lucy Anne Wilson, Mari Herigstad

**Affiliations:** Biomolecular Sciences Research Centre, Sheffield Hallam University

## Abstract

Individuals are routinely exposed to low-level carbon monoxide (CO), by factors such as ambient pollution and tobacco smoking. It is known that inhalation of high levels of CO have a detrimental impact on cognitive function. This study sought to investigate the impact of low-level CO exposure on central nervous system cognitive processing speed, using Blood Oxygen Level Dependant (BOLD) functional Magnetic Resonance Imaging (fMRI). The effects of low-level CO (raised up to 6ppm in exhaled air) on reaction times and fMRI activation maps were measured in healthy non-smoking participants. Participants received BOLD fMRI scans on two separate occasions (air and CO intervention days) and were scanned during the performance of a simple reaction time task. Results showed mean activation in cerebellum and motor cortex for all conditions. A significant reduction in BOLD response in the right temporal gyrus was found following CO inhalation, compared to the air control. Reaction times were significantly slower after CO exposure on the CO experimental day, but did not significantly change on the air control experimental day. This suggest that even low-level CO may impact both behavioural and BOLD fMRI outcomes.

## INTRODUCTION

Most individuals are exposed to ambient levels of CO every day (Penney et al., 2010). Government strategies detail 9 parts per million (PPM) as acceptable exposure (Department for Environment Food & Rural Affairs, 2019) however lower levels than this threshold have been linked with cardiovascular diseases, such as stroke (Lee et al., 2014; Maheswaran et al, 2005; Hedblad et al, 2005) and heart failure (Shah et al, 2013), as well as altered cardiac embryonic development (Matias et al., 2024). Cardiovascular diseases remain a key risk factor for neurological impairment, and the ‘acceptable’ levels of CO have been associated with vascular dementia onset (Chang et al, 2014).

Whilst several studies focus on effects of acute, high levels of CO in human populations exists (Eichhorn et al., 2018), there is limited research available on low-level CO inhalation, with most evidence emerging from studies that are observational in nature (Yoon et al, 2020). Whilst a few epidemiological studies have identified an association between low-level CO and disease risk (Lee et al., 2014; Maheswaran et al, 2005; Hedblad et al, 2005) CO in air pollution is part of a cocktail of potentially damaging molecules, and it is difficult to disentangle the impact of CO alone on health. Furthermore, the cognitive effects of relatively low-level CO exposure are also as of yet poorly understood.

A recent study used functional Magnetic Resonance Imaging (fMRI) to assess breathing, motor and visual responses to low-level CO (3ppm) and identified that this low level altered fMRI signal (Bendell et al, 2019). fMRI is widely regarded in the neuroimaging community as a fundamental measurement of neurological structure and performance (Zhang et al., 2019). Blood oxygen level dependent (BOLD) fMRI is frequently used to non-invasively measure neural activity, through the interpretation of mechanisms that impact BOLD signal such as region-specific metabolism requirements and complex vascular relationships (Gauthier & Fan, 2019). BOLD fMRI contrast images are developed in response to the differing magnetic properties of oxygenated vs deoxygenated haemoglobin, whereby oxygenation increases signal intensity (Zhang et al., 2019). As such, increasing neurological demand in a brain region simultaneously leads to an increase in local cerebral blood flow (CBF) and a relative change in oxygenated and deoxygenated blood, which forms the basis of the BOLD signal (Drew, 2019).

Numerous tasks have been conducted in fMRI research, and reaction time tasks are among the most established (Yaple et al., 2019). Existing studies have identified strong similarities between participant responses when conducted inside and out of the scanner, suggesting that in-scanner performance can be regarded as largely reliable (Koten et al, 2013).

Furthermore, reaction time task results have been shown to be effective measures of BOLD response across a diverse age range, particularly when interested in the cerebellum region (Yaple et al., 2019). In this study, a reaction time task was employed to explore how BOLD response may be affected by low-level CO, during performance.

## MATERIALS AND METHODS

### Participants

A total of 12 never-smokers were recruited to take part in the study. Participants were in general good health, on hormonal contraceptive if female and provided written, informed consent prior to enrolment. Smokers characteristically exhibit elevated levels of COHB (Pan et al, 2021) and thus were excluded from the recruitment population. Additional exclusion criteria were MRI contraindications (e.g., pacemaker), pregnancy and any background of neuro/cardiovascular or respiratory condition or impairment. The principles of the Declaration of Helsinki were adhered to throughout the study. Parts of this work has been reported elsewhere (Bendell et al., 2019).

### Protocol

Prior to study commencement, participants attended an introductory visit, where details of relevant histories and State-Trait Anxiety Inventory (STAI, Spielberger et al, 1983) questionnaires were completed to obtain baseline indications of anxiety or distress (Green et al, 2017). Participants were provided opportunity to familiarise themselves with the custom-made breathing apparatus, consisting of a mouthpiece and nose clip to prevent nasal air entry. Once comfortable and stable respirations had been observed for 5 minutes, low volume CO was introduced to the breathing circuit to mix with inspired air. This activity was concealed from the participant and then an additional 5 minutes of respirations were recorded. Physiological monitoring was maintained throughout the intervention, including oxygen saturation, heart rate monitoring and ECG. The Micro+ Smokelyzer kit (Intermedical LtD, Kent) was employed to measure expired CO levels before, immediately after and 10 minutes after the breathing test was complete.

Participants underwent MRI scans across two separate days and were required to complete the anxiety section of the STAI on arrival to the department and within 15 minutes of finishing the experiment each day. A variety of tasks were completed whilst in the scanner, including a reaction time task (other tasks reported elsewhere, see Bendell et al. 2019). Training was provided on the completion of all tasks before entering the scanner on both scan days, to ensure task proficiency when alone in the scanner and participant comfort.

The reaction time task involved the pressing of a button with the individual’s right hand, as soon as a red dot flashed up on screen. Participants were required to perform this action as fast as possible for a total of 24 dot appearances, shown at random intervals, and their reaction times were recorded. Individuals completed all tasks once before the breathing intervention began (baseline measurement) and once more after the gas mixture was introduced (post-intervention). Participants inhaled the gas mixture inside the scanner. Participants were block randomised to receive an air intervention on one scan day and a CO intervention on the other.

Participants were unaware of which order they were receiving the intervention but were aware that CO would be introduced into the breathing circuit at some point. Participants were asked at the end of each scan day if they could guess which intervention had been given to them (air or CO) but were unable to reliably tell the difference. Expired CO levels were obtained prior to the first scan, immediately after the final scan (approximately 20 minutes post intervention) and again 10 minutes after the post-exposure scan (approximately 30 minutes post intervention).

### Data Collection and Analysis

A Siemens 3 Tesla TIM-Trio scanner with head coil was used to obtain two fMRI scans per participant (BOLD echo-planar image acquisition, time repetition (TR) = 3000ms, time echo (TE) = 30ms, field-of-view = 192 x 192mm, voxel-size=3x3x3mm, 45 slices) on each scan day. Obtained scans were structurally registered against a T1-weighted brain scan (MPRAGE, TR=2040ms, TE=4.7ms, flip angle=8°, voxel-size=1x1x1mm; Bendell et al, 2019). All physiological measurements were continually obtained at 50 Hz using via PowerLab 16/35 using LabChart, which facilitated simultaneous biological monitoring and scan acquisition with reasonable accuracy (Mahdiani et al, 2015).

Scan data was processed using FEAT (FMRI Expert Analysis Tool) version 6.0 within FSL (Oxford Centre for Functional Magnetic Resonance Imaging of the Brain (FMRIB) Software Library). A cluster-defining Z threshold was set to 3.1 to test for significance at the alpha level of 0.05. Motion correction in the form of MCFLIRT was applied, and further prestatistical processing included spatial smoothing with a full-width-half-maximum Gaussian kernel (5mm) and high-pass temporal filtering.

First-level analysis consisted of running each participant’s scan (4 in total) with a model comprising of the individual reaction times for that condition. Reaction times were introduced using a 3-column design, with the time interval of how far into the scan the dot was presented in seconds (s, column 1), how long the dot was shown for (0.16s each time, column 2) and the speed at which the participant responded in seconds (column 3). EVs were convoluted using a gamma waveform to approximate the canonical haemodynamic response (Lindquist et al, 2012), and physiological noise was regressed out using a physiological noise modelling tool. A 6-s haemodynamic delay was assumed. Lower level analyses were combined for each participant using a second higher level fixed effects analysis, comparing baseline and post intervention scans. Group analysis calculated group means for each of the four scan sessions (pre and post air/CO), and a repeated measures full group analysis was done to compare air and CO experimental days. A hierarchical approach to effects modelling was adopted (Woolrich et al, 2004). For higher-level group analyses, a mixed-effects model, FLAME (FMRIB’s Local Analysis of Mixed Effects, Woolrich et al, 2004) was adopted.

Physiological data (PO2, PCO2, respiratory rate, heart rate) was processed using custom Matlab scripts (Garry et al, 2016) and analysed individually using SPSS (IBM, 2019). SPSS was also used to assess individual reaction times of participants and visualise outliers through boxplot generation (Parke, 2013). Reaction times were plotted using pre and post intervention scan data, and outlier information and mean values were calculated. Reaction times, anxiety scores and CO exhaled levels were compared using individual Student’s T-tests.

## RESULTS

Ten participants yielded usable imaging data for analysis. The two participants that did not have suitable imaging data (due to missing reaction time data) were excluded from all further analysis. Table 1 shows demographic data for the ten participants with usable imaging data.

**Table 1.** Participant demographic and behavioural data.

|  | Preliminary visit | MRI visit (CO) | MRI visit (Air) |
| --- | --- | --- | --- |
| Sex (F/M) | 6/4 | 6/4 | 6/4 |
| Age (years) | 25.3 +/-4.8 | 25.3 +/-4.8 | 25.3 +/-4.8 |
| BMI (kg/m<sup>2</sup>) | 24.0 +/- 3.2 | 24.0 +/- 3.2 | 24.0 +/- 3.2 |
| State anxiety score | 3.1 (8.6) [21-55] | 27.0 (4.3) [21-35] | 28.2 (4.4) [23-37] |
| RT change (post>pre, s) | N/A | +9.2 | +13.9 |
| *Mean (SD). Range included for state scores. BMI: body mass index; RT: reaction time.* |  |  |  |

All participants showed a significant increase in expired CO level after the CO gas inhalation (baseline 3.0ppm+/-1.1, post CO 5.8ppm+/-0.8, p<0.001), but not for air (baseline 3.2ppm+/-0.9, post air 2.8ppm+/-0.8, NS). Group comparisons showed a significant rise in CO in the CO protocol (2.8ppm+/-0.9) compared to air (-0.4ppm+/-0.7, p<0.001). Participants showed no significant difference in anxiety between the air and CO protocols (p=0.08). Participants showed no difference in any physiological parameters between air and CO protocols.

Participants demonstrated slower reaction times on average in their post-exposure scans (post air = 0.374s+/-0.043; post CO = 0.388s+/-0.042), compared to baseline scans (air = 0.365s+/-0.041, p=0.06 (NS); CO = 0.374s+/-0.040, p=0.03), regardless of experimental protocol. Reaction times exhibited post CO were slowest overall, and were significantly slower than baseline reaction times on the same day (p=0.03). This was not seen in the air protocol (figure 2). There were no outliers amongst the participants for any of the tasks.

**Figure 1.**
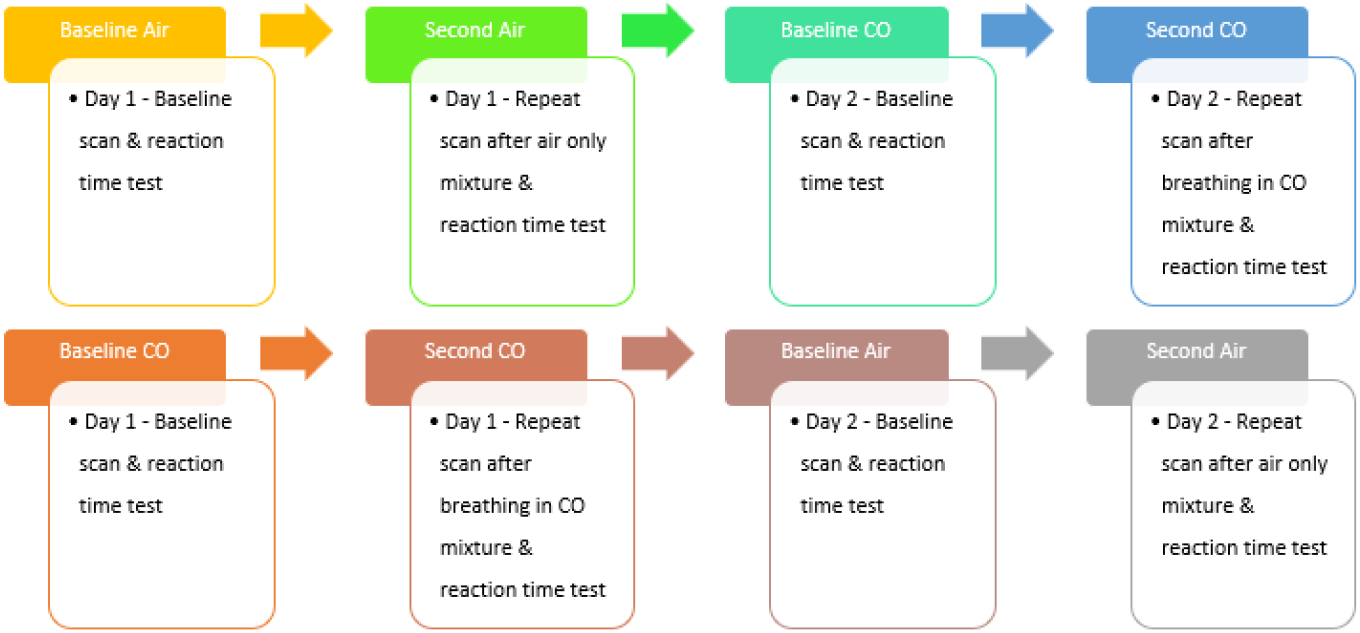
Schematic of the experimental paradigm. Half of participants were given air on their first experimental day and CO on the second (top), and half were given CO on their first experimental day and air on the second (bottom).

**Figure 2.**
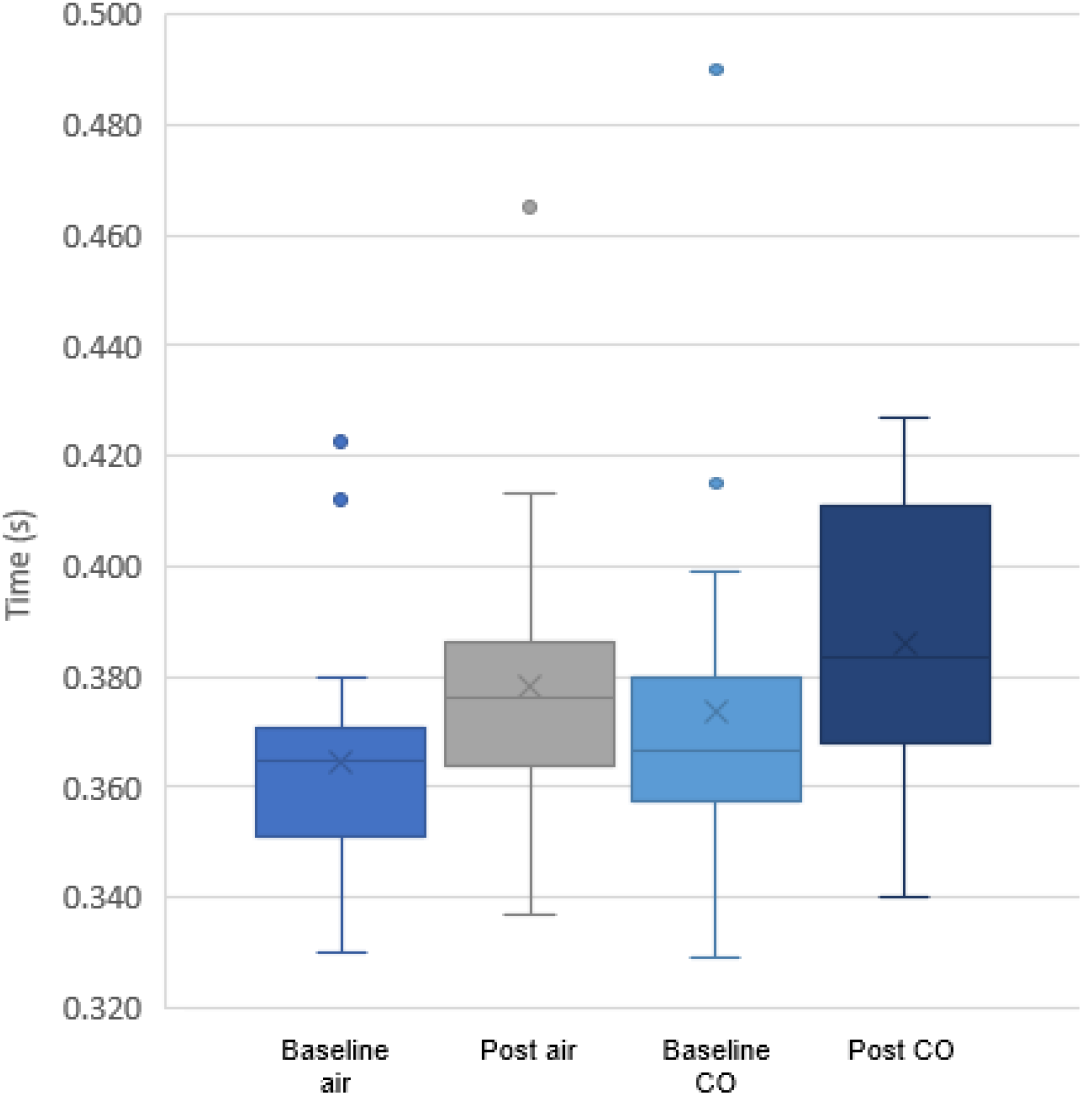
Reaction time data. Participants demonstrated slower reaction times in their post-intervention scans compared to baseline scans on each day. Reaction times were significantly slower following the gas inhalation in the CO protocol, but not in the air protocol..

Group mean analysis showed uniform activation in the cerebellum for CO and air protocols, though activation decreased following CO (figure 3). The motor cortex also exhibited consistent voxels of activation, although signal was reduced post-CO (figure 4). The post air scan furthermore showed activation in the somatosensory cortex, thalamus, visual cortex, and middle temporal gyrus. The baseline CO scan showed additional activation in the cingulate gyrus, premotor cortex, thalamus and anterior cingulate gyrus. The post CO scan showed activation in the corticospinal tract (right) and the primary somatosensory cortex.

**Figure 3.**
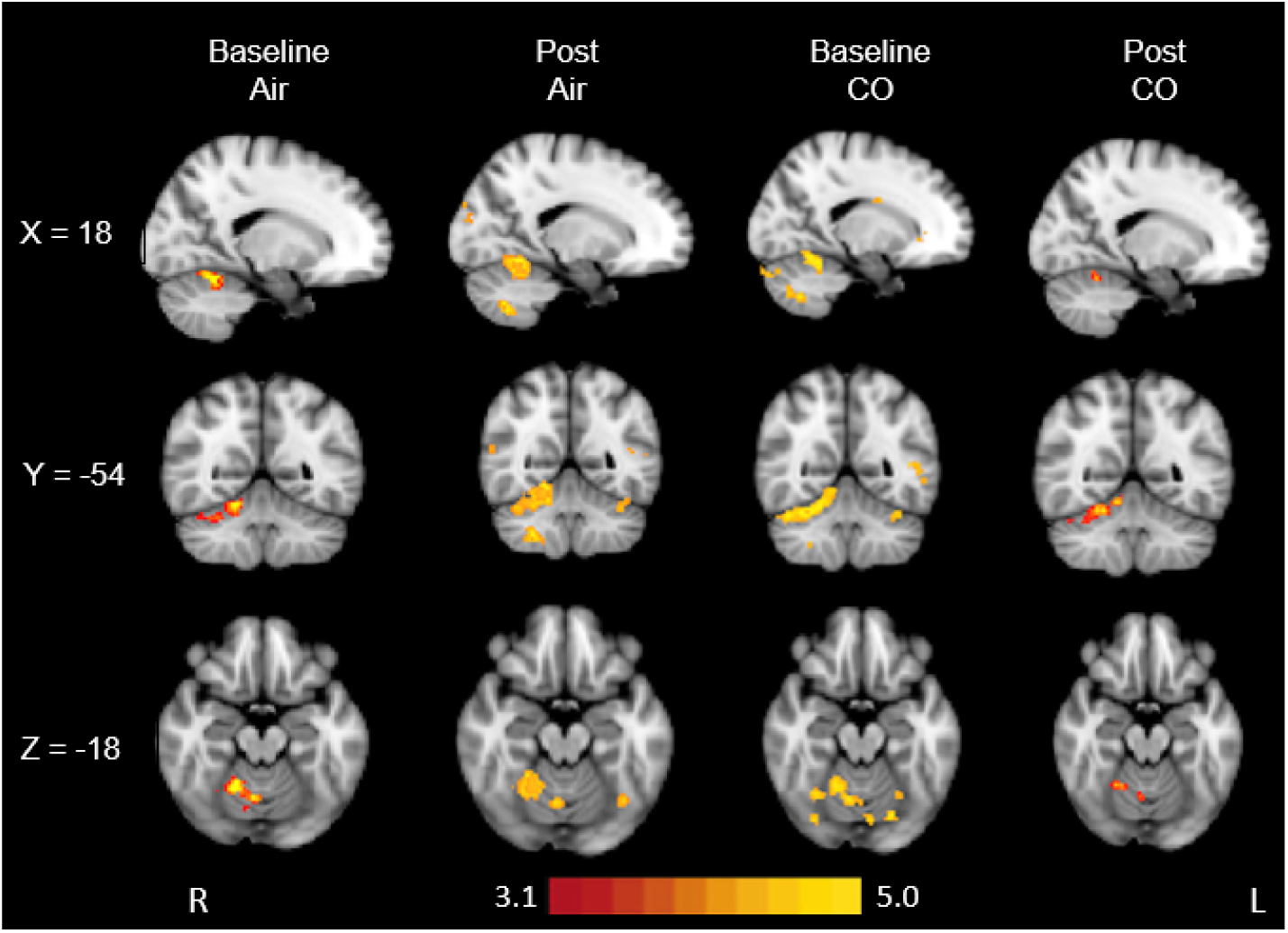
Mean cerebellar activation for all scans, whole brain analysis. Maps are statistical maps (Z scores) superimposed on a standard (MNI) T2 2mm brain. Significant regions are displayed with a threshold of Z□> □3.1 with a cluster probability threshold of p□< □0.05 (corrected for multiple comparisons).

**Figure 4.**
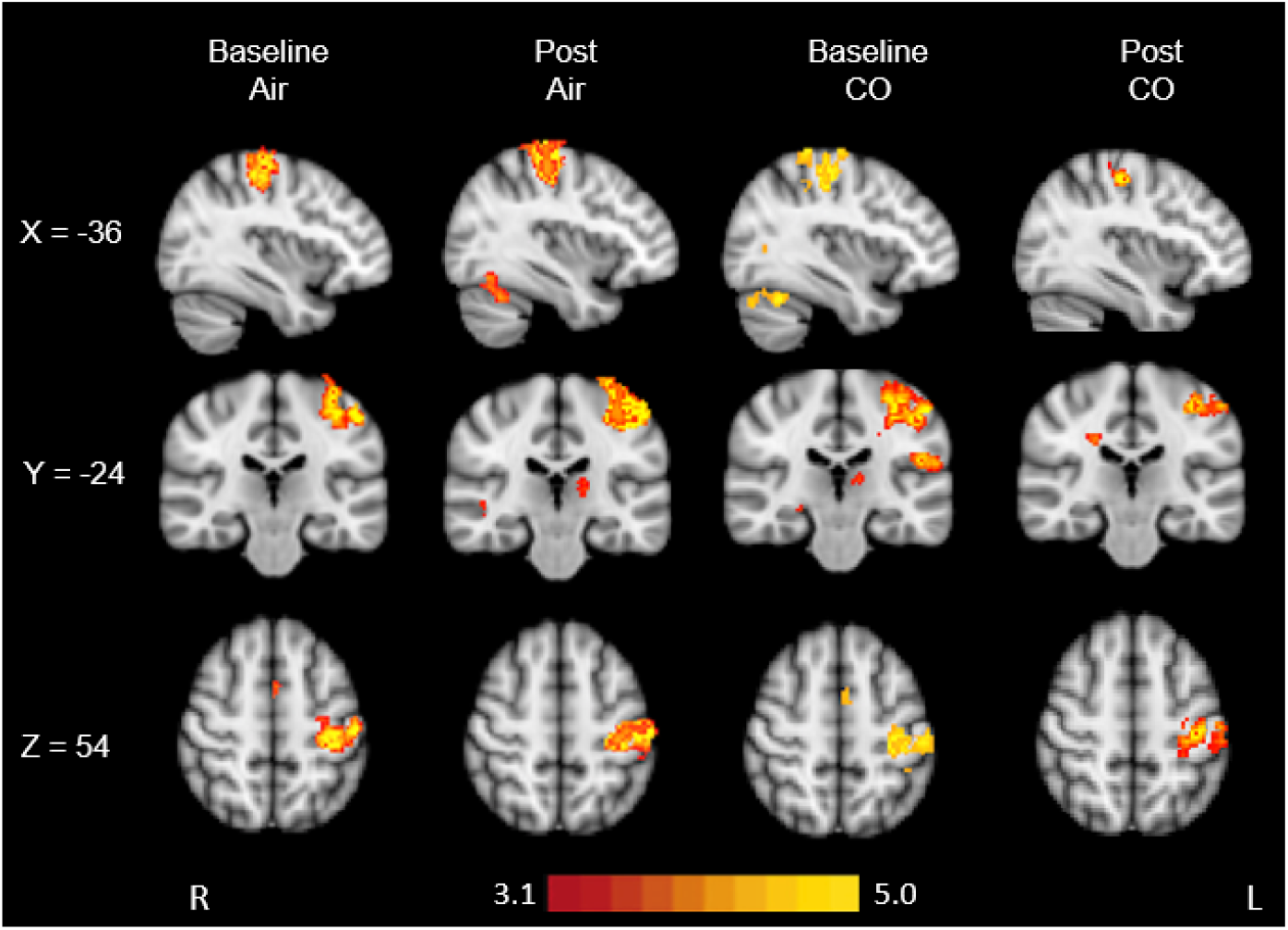
Mean motor cortex activation for all scans, whole brain analysis. Maps are statistical maps (Z scores) superimposed on a standard (MNI) T2 2mm brain. Significant regions are displayed with a threshold of Z□> □3.1 with a cluster probability threshold of p□< □0.05 (corrected for multiple comparisons).

Comparison between the air and CO showed significantly reduced BOLD responses after CO exposure compared with air exposure in the middle temporal gyrus (figure 5), meaning that on the day the participants inhaled CO, the BOLD response was reduced in the post-inhalation scan (CO(post>pre)) compared to the day the participants inhaled air (Air(post>pre)). BOLD response was in no area increased following CO compared to BOLD response following Air.

**Figure 5.**
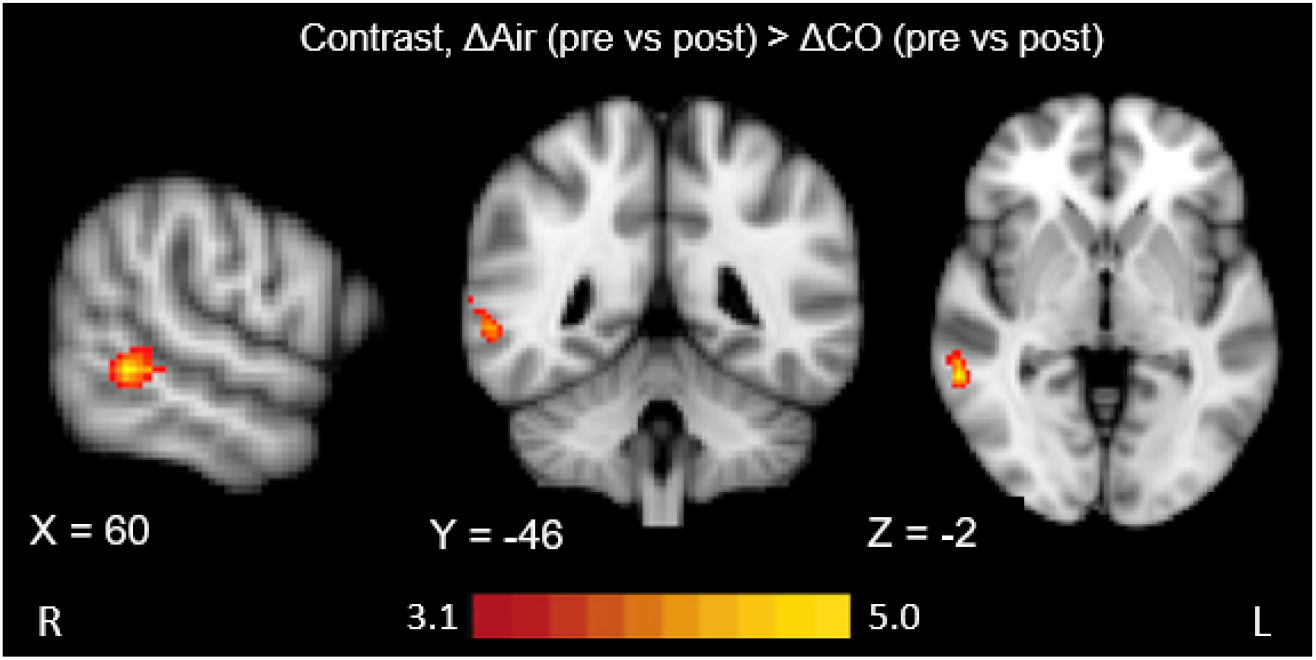
CO impact on BOLD fMRI response, whole brain analysis, group comparison. Red-yellow scale indicates where cerebral activation was lower post-CO compared to post-air. Maps are statistical maps (Z scores) superimposed on a standard (MNI) T2 2mm brain. Significant regions are displayed with a threshold of Z□> □3.1 with a cluster probability threshold of p□< □0.05 (corrected for multiple comparisons).Harvard cortical and subcortical atlases (FSL) determined that reduction in BOLD response occurred in the middle temporal gyrus.

## DISCUSSION

Here we show that inhaling a low amount of CO (raising expired CO levels with approximately 3ppm to 6ppm) is sufficient to alter cerebral BOLD response during a reaction time task in a group of non-smokers. This was paired with a reduced reaction time following the CO inhalation. These findings suggest that heightened carboxyhaemoglobin (COHb) may be a non-negligible confound, particularly in BOLD fMRI studies, and it highlights how even low doses of CO may impact function. Though we cannot determine the mechanism underlying this altered BOLD response, potential contributing factors are outlined below, alongside recommendations for future research.

Participants in this study undertook repeated reaction time tests, selected due to reported high validity and test-retest reliability when conducted in previous fMRI studies (Koten et al, 2013). Reaction time data was collated for all participants and grouped by scan type, before averaging the reaction speed across individuals for each of the four conditions. We observed good adherence to the task, with few events missed for all protocols. Standard deviations were small, suggesting that reaction times were consistent between participants, and response times were within expected ranges.

We observed slower reaction times in both protocols, suggesting perhaps an element of e.g. tiredness or similar in the latter half of the scan. As reaction time tests have the potential to reveal inter-individual differences in e.g. anxiety (Riedel et al, 2016) which could in turn influence BOLD response or slow reaction times (Ford et al, 2018), we assessed state anxiety pre and post scan for both protocols. There was no change in state anxiety, suggesting that the change in reaction times should not be attributed to feelings of participant apprehension.

The slower reaction time was furthermore not significant in the control air protocol, but did reach significance in the CO protocol (figure 2). This suggests that there was an impact of CO on processing speed in our participants. Indeed, CO in higher doses is known to affect cognition, with attention, immediate memory, and processing speed being consistently affected (Watt et al, 2018).

Nevertheless, future research might employ different tasks to ensure that participant engagement is maintained, and fatigue is minimised wherever possible. The effect of acute high dose CO and decline in cognitive processing abilities is well documented (Pages et al, 2014; Chen et al, 2013), but these findings highlight how neurological behaviour may be influenced by significantly lower levels of CO exposure.

Mean activations were examined for the CO and air protocols, and a strong BOLD response was visible in the cerebellum and motor cortex for all scan types (figure 3, figure 4). Reaction time tests have been proven to elicit multi-region brain activation (Yarkoni et al, 2009), thus maximising the opportunity for changes in BOLD response to be witnessed. The large, uniform clusters of activation generated in both the cerebellum and motor cortex reinforce the replicability of reaction time tasks in fMRI BOLD studies (Bossier et al, 2020).

The cerebellum is responsible for movement execution and coordination (Dahlberg et al, 2020, Hwang et al, 2009), and also attention and sensory processing (Phillips et al, 2015). Cerebellar activation during the performance of the reaction time tasks was thus expected. Motor tasks are initiated and maintained by a complex series of cerebral connections, largely controlled by the motor cortex (Khushu et al, 2001). As anticipated, the group mean analyses revealed large scale similar activation across all scan types in the left side motor cortex, consistent with the participants pushing a button with their right hand during the scan. Additional BOLD response is seen in the supplementary motor cortex in both baseline scans, although not in the post-exposure scans. This suggests that task repetition may have impacted cerebral blood flow in this area, perhaps in response to increased cerebrovascular efficiency (Toma et al, 1999). However, whilst the activation in the motor cortex are similar across the scans (figure 4), there is an observable reduction in BOLD response in the post CO condition. This reduction, unlike the one in the supplementary motor cortex activation, cannot reliably be explained by task repetition phenomena (Mazzetto-Betti et al, 2011), due to consistent activation in both the pre and post air scans. While it is possible that the CO exposure caused this trend towards signal reduction in line with the findings in Bendell et al. (2019), given that neither of the above trends reached significance, we cannot draw any such conclusions.

Thalamus activation was detected in the air scans but diminished in the post-CO scan type. Research indicates that thalamus impact following CO inhalation is rare (Abdelhedi et al, 2013; Lo et al, 2007). The thalamus has a core role in the relaying of sensory information, including motor control responses, and so would naturally be involved in this task (Torrico & Munakomi, 2022). This could describe how with reduced thalamus activity, reaction times lengthened, though it is unclear whether the presence of CO was a contingent or coincidence.

Studies have identified a link between increased reaction times and reduced BOLD response (Firbank et al., 2017), although the precise underpinning factors are ambiguous. Potentially this could be caused by a change in haemodynamic behaviours and as such tissues are not sufficiently perfused to perform cognitive tasks efficiently. Alternatively, if cognitive processing abilities have been affected, targeting blood flow may be reduced and as such the visible BOLD response is limited. Although it is uncertain, the diminishing BOLD response viewed in the post-CO scan set could be attributed to either one of these factors.

A statistically significant reduction in BOLD response following CO (figure 5) was evident at group level in the superior and middle temporal gyri, located in the temporal lobe (Patel & Fowler, 2021). Cerebral damage following CO toxicity displays a predilection for the temporal lobe (Zhang et al, 2019; Chen et al, 2013), and so is a region of interest for the effects of low-level exposure. The superior and middle gyri regions are commonly associated with sensory processing and semantic memory (Onitsuka et al, 2009). These areas were likely involved as the participants observed visual dot flashes and recalled the need to press a button in response. The superior temporal gyrus has further been linked to the processing of eye movements (Stigler & McDougle, 2013) and may have been implicated through the perception of dot flashes. Research into neurological degenerative diseases have described a link between oxidative stress and temporal gyri abnormalities (Lin et al, 2015), suggesting that temporal regions may be vulnerable to the effects of CO and the associated physiological chain-reactions.

CO exhibits vasodilatory properties and may modify levels of nitric oxide (NO) in the bloodstream (Choi and Kim, 2021). Research into the purposeful utilisation of low-level CO as a vasodilator in gestational hypertension has been presented in rodent models (Venditti et al, 2014), though it’s relationship with cerebral responses is sorely misunderstood (Townsend and Maynard, 2002). Our findings show a reduction in BOLD signal following CO inhalation, in line with Bendell et al. (2019), and may be a consequence of increased baseline vasodilation. If the cerebral vasculature is pre-dilated, this could leave less potential for further dilation, and thus reduce the task-related BOLD signal as this depends upon a comparison with baseline signal. However, as the altered BOLD signal was linked to a measurable difference in function, namely reaction time following CO exposure, it is possible that the impact of low-level CO on the human brain goes beyond a simple vasodilatory response. Rather, it may be that CO influences both cerebral vasculature (mainly in the form of modifying vasodilation through its impact on NO) and on the neuronal function. The underpinning mechanism for this remains unknown.

### Limitations

Whilst reaction time tasks are regarded a reliable and valuable measure for cognitive responses, the threat of attention loss and variability in speed between participants threatens its validity (Riedel et al, 2016). While it is likely that any habituation to the respiratory breathing apparatus was minimal (Hayen et al, 2015), habituation could have impacted the reaction time task to some degree. A non-significant slowing of response times was found in the air protocol which could suggest some level of attention loss or fatigue, but this is unlikely to fully explain the significantly greater increase in response times following CO. Nevertheless, future works should consider an enhanced, multi-level approach to participant reaction time testing, perhaps by combining the n-back paradigm to increase complexity for volunteers.

## CONCLUSION

In summary, this study indicates that even low levels of CO are sufficient to alter cerebral vascular activity during task performance as well as performance output (in the form of reaction times). Further research into potential underpinning mechanisms should be explored in future work to aid generalisability.

## ACKNOWLEDGEMENTS

This study was funded by the Oxford Brookes University Central Research Fund. MH and LW are supported by the Carbon Monoxide Research Trust.

